# OptoRheo: Simultaneous *in situ* micro-mechanical sensing and imaging of live 3D biological systems

**DOI:** 10.1101/2022.04.21.489042

**Authors:** Tania Mendonca, Katarzyna Lis-Slimak, Andrew B. Matheson, Matthew G. Smith, Akosua B. Anane-Adjei, Jennifer C. Ashworth, Robert Cavanagh, Lynn Paterson, Paul A. Dalgarno, Cameron Alexander, Manlio Tassieri, Catherine L. R. Merry, Amanda J. Wright

## Abstract

Biomechanical cues from the extracellular matrix (ECM) are essential for directing many cellular processes, from normal development and repair, to disease progression. To better understand cell-matrix interactions, we have developed a new instrument named ‘OptoRheo’ that combines light sheet fluorescence microscopy with particle tracking microrheology. OptoRheo lets us image cells in 3D as they proliferate over several days while simultaneously sensing the mechanical properties of the surrounding extracellular and pericellular matrix at a sub-cellular length scale. OptoRheo can be used in two operational modalities (with and without an optical trap) to extend the dynamic range of microrheology measurements. We corroborated this by characterising the ECM surrounding live breast cancer cells in two distinct culture systems, cell clusters in 3D hydrogels and spheroids in suspension culture. This cutting-edge instrument will transform the exploration of drug transport through complex cell culture matrices and optimise the design of the next-generation of disease models.

## Introduction

Cells sense and respond to the mechanical properties of the extracellular matrix (ECM) at a cellular length scale, using traction forces to probe stiffness^1^, steer migration^2,3^ and influence cell fate^4^. Simultaneously, the ECM is continuously remodelled by cells as they exert these traction forces^5^ during cell migration and morphological re-arrangement^6^. Anomalies in the mechanical properties of the ECM play significant roles in the development of pathologies such as cancer^7^ and fibrosis^8^, often establishing barriers to therapeutic intervention^9^. Modelling and understanding cellular influence on ECM biomechanics is challenging given the wide range of mechanical environments experienced in health and disease. In healthy tissues, the elastic modulus has been reported to range from tens of Pa (e.g., brain, lung) to well above 10 KPa (e.g., skeletal muscle, bone), with disease states such as cancer and fibrosis showing a significant change in stiffness (e.g., from 800 Pa for normal breast to more than 4 KPa in breast cancer)^10^. Moreover, the full mechanical characterisation of the ECM also contains a viscous component that may influence cell behaviour^11^. The recent development of engineered hydrogels with tuneable mechanical properties^12,13^ have made it possible to recreate elements of the ECM micro-architecture *in vitro* and reveal the influence of ECM viscoelasticity on cell processes^14,15^. Despite these advances and their importance, the mechanistic processes of cell-matrix interactions remain poorly understood. For instance, do cells ‘prime’ their local environment prior to migrating or do they exploit existing weaknesses in the ECM and migrate accordingly? These unanswered questions call for minimally invasive optical approaches to monitor changes in the microscopic mechanical properties of the ECM, *in situ* and in real time, local to proliferating cells over many days.

To address this aim, OptoRheo combines three different microscopy techniques, light sheet microscopy, multiplane microscopy and optical trapping into a single instrument. This approach enables live fluorescence imaging deep in 3D cell cultures and microrheology measurements of the ECM within the same region of interest, local to and far from the cells. 3D fluorescence imaging is achieved using a new version of reflected light sheet fluorescence microscopy (LSFM)^16–19^ built on a commercial inverted microscope body to image hundreds of microns deep from the coverslip, within live 3D cell cultures and with sub-cellular resolution. The sample is kept completely stationary during z-scanning, with no perturbation or contamination risk from dipping lenses, both crucial for ensuring that observation does not influence the mechanical or biological properties of the sample. This novel configuration allows for delicate samples such as hydrogel scaffolds to be imaged simply in off-the-shelf chambered coverslips. To extract the viscoelastic properties of the ECM, OptoRheo tracks the thermally driven Brownian motion of micron-sized beads, acting as rheological probes across a wide time-window. The inert bead probes can be embedded in the hydrogel during encapsulation^6,20–22^ or even internalised into cells^23^ to probe intracellular viscoelasticity. In suspension cultures, an optical trap can be used to hold the probe in the field of view within the cells’ microniche during the measurement extending the range of materials the instrument can characterise^24^. Finally, OptoRheo incorporates optional multiplane imaging that can be used to extend microrheology to 3D in a configuration similar to the one developed for OptiMuM^25^, to achieve a full 3D characterisation of the extracellular microenvironment.

To highlight the capability of OptoRheo, we present data obtained from the analysis of two systems seeded with human-derived MCF-7 and/or MDA-MB-231 breast cancer cells, either (I) encapsulated as clusters in 3D hydrogels or (II) as spheroids maintained in suspension culture. In the case of hydrogel scaffolds, matrix stiffness was measured using passive particle tracking microrheology without the use of an optical trap or multiplane imaging, whereas in the case of suspension cultures, the optical trap was implemented along with multiplane imaging. Imaging and microrheology were performed sequentially at multiple regions within the samples at depths of 150 μm - 400 μm from the coverslip. In the case of the hydrogels, the samples were monitored over three days to reveal microscale variations in the elastic properties of the ECM near to and away from cells. When studying spheroids in suspension the optical trap was used to place and hold the probes in user-defined locations and extract the relative viscosity of the media near the spheroids. Notably, in both the cases our measurements were found to be sensitive to local spatio-temporal variations in the biomechanical properties of the culture medium. As demonstrated in this study, our multimodal and minimally invasive approach opens a wide range of future opportunities for physiologically relevant, long-time course investigations and time-lapse videos of cell-ECM interactions in fragile live cell culture samples. We anticipate that this approach will be applied to increasingly complex and relevant *in vitro* models, providing an essential insight into the previously opaque mechanistic control of cell behaviour by the ECM in health and disease.

## Results

### 3D imaging deep in live cell cultures

A schematic representation of OptoRheo can be seen in Figure 1. 3D light sheet fluorescence microscopy (LSFM) is achieved by projecting a thin, planar excitation beam limited to the detection plane of the microscope and collecting the emitted fluorescence at a 90° angle to the illumination plane^26,27^. For deep imaging of live 3D cell cultures, the light sheet illumination was introduced using a 10 mm 90:10 (Reflectance: Transmittance) beam splitter cube placed in the sample chamber prior to casting the gel alongside it (Figs 1, 2 and S1). This LSFM approach has multiple advantages; deep and fast fluorescence imaging with low phototoxicity^28^ and minimal sample perturbation during imaging, while being cost-effective and modular. The glass beam splitter cube can be sterilized for reuse and placed either inside or outside the sample chamber to provide flexibility to adapt to different experimental conditions.

**Figure 1:**
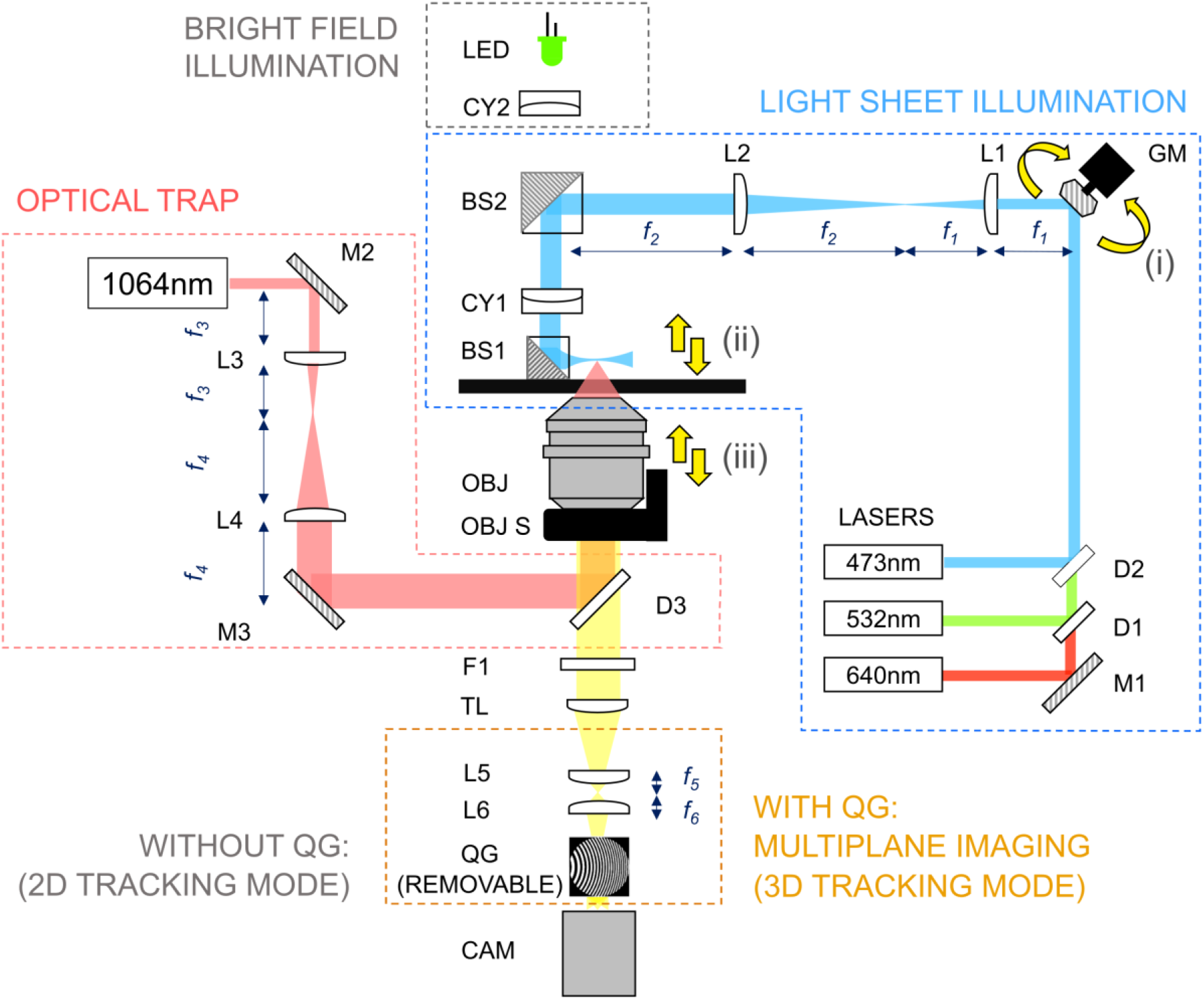
A schematic representation of the OptoRheo instrument. Components: Lasers – 473 nm, 532 nm and 640 nm lasers provide the light sheet illumination while a 1064 nm laser is used for optical trapping. M1- M3 – mirrors; D1-D3 – dichroic mirrors; L1-L6 – achromatic doublets, L1 and L2 form a 8.3 x beam expander and are part of a 4f system with the galvanometer mirror (GM) and a beam splitter (BS2); CY1, CY2 – cylindrical lenses; BS1, BS2 – beam splitter cubes; OBJ – objective lens; OBJ S – piezoelectric objective scanner; F1– fluorescence emission filter; TL – tube lens, QG – quadratic gratings and CAM – camera. Yellow arrows indicate synchronised motion of (i) the galvanometer mirror, (ii) the projected light sheet and (iii) the objective lens (using a piezoelectric objective scanner). The quadratic gratings slide in and out of the optical path to enable 2D and 3D particle tracking. The quadratic gratings are removed for the LSFM imaging.

**Figure 2:**
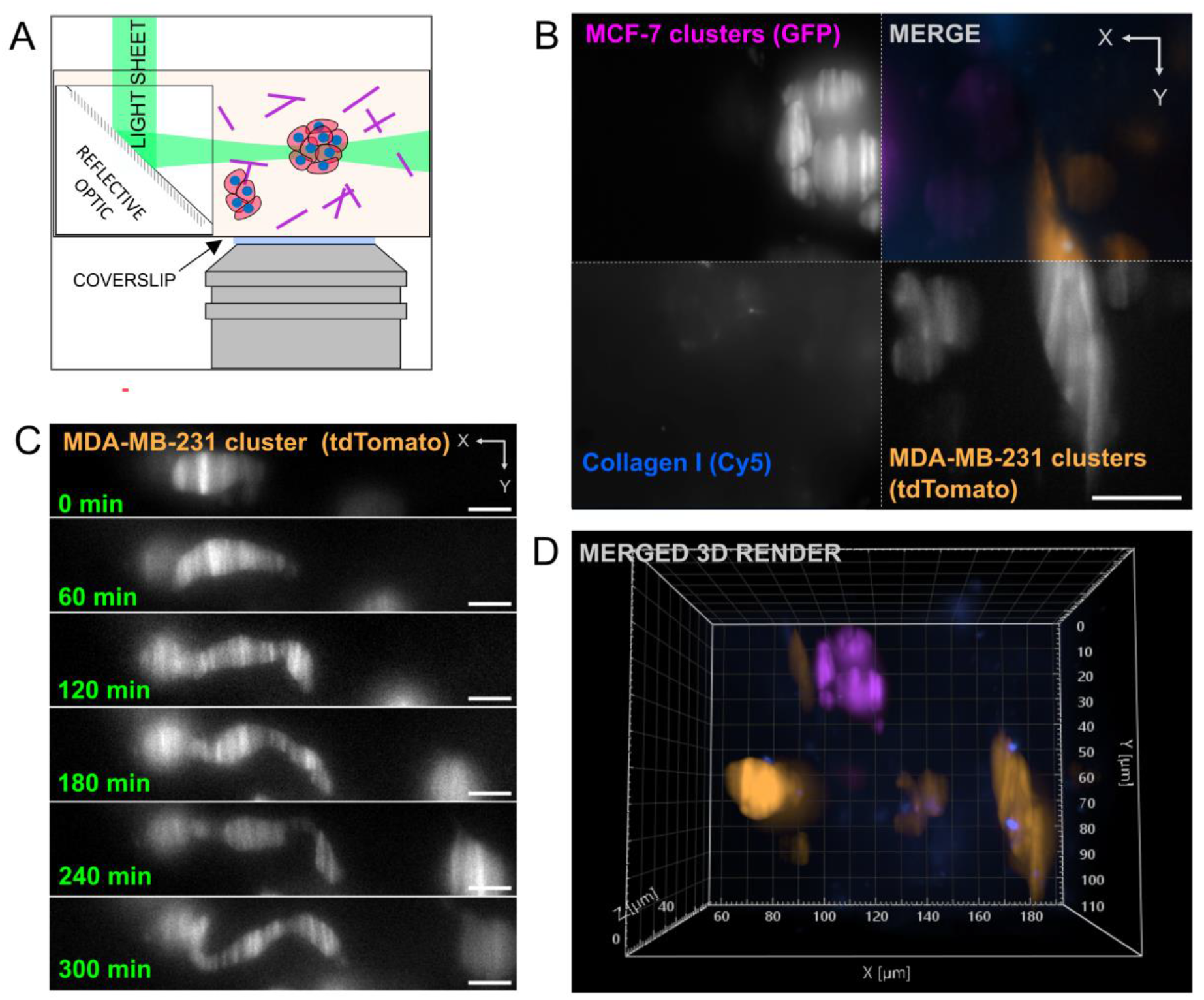
Light sheet fluorescence microscopy (LSFM) with OptoRheo. A. Schematic of light sheet microscopy on the OptoRheo. Sample consists of cell clusters (represented by red circles with blue centres) and collagen (purple lines). B. Multicolour imaging of a co-culture of MCF-7 (top left corner of montage) and MDA-MB-231 (bottom right corner) in hydrogel supplemented with Cy-5 labelled collagen I (bottom right corner) on the OptoRheo with three colour channels. Images in montage are maximum intensity projections. Scale bar = 20 μm. C. Single planes from time lapse movie (supplementary video VS1) taken on the OptoRheo of a MDA-MB-231 cell changing morphology. Scale bar = 10 μm. D. 3D rendering of the same region of interest in panel B.

Unlike some other prism or mirror based LSFM solutions^16,29,30^, 3D image generation was achieved here by scanning the light sheet and not the sample. Notably, this allows the sample to be kept stationary and undisturbed throughout data collection, which is essential for imaging delicate samples prepared in soft hydrogels (Fig 2A) or liquid suspension media over multiple days. The light sheet itself was generated using a cylindrical lens, the properties of which set the thickness of the light sheet and influence the axial resolution and optical sectioning capabilities of the microscope. In the presented configuration, the measured axial resolution of the detection optics of the LSFM on OptoRheo was 1.09 μm for λ_ex_\ λ_em_ = 532 nm \ 580 nm which agrees closely with theory (1.1 μm) (Fig S2).

A scanning galvanometer mirror was placed conjugate to the cylindrical lens using a 4f system, so that tilting the galvanometer mirror translated to a Z-shift in position of the light sheet at the sample (Fig 1). Acquiring Z-stacks involved synchronisation of the galvanometer mirror with a piezoelectric objective scanner that moved the objective lens, ensuring that the illumination and imaging planes remained co-aligned and synchronised throughout the scan. The light sheet remained at optimal thickness (~3 μm for all three colour channels, see Methods) over a field of view of ~100 μm. However, image tiling could be achieved within a region 4 - 6 mm from the beam splitter cube to increase the field of view. This required shifting the light sheet focus laterally by moving the position of the cylindrical lens. The shifted position of the beam splitter was compensated by tilting the galvanometer mirror to image the new focal position.

The current configuration of OptoRheo uses a 60x objective lens with a 1.5 mm working distance and a numerical aperture (NA) of 1.1, selected to image deep into a sample, but with a high enough NA for optical trapping. For the data presented in this work, z-scans were typically recorded 150 μm - 400 μm from the coverslip (Fig 2B). To extend the field of view for larger objects, LSFM images could be tiled and stitched together as detailed above. The multiplane grating breaks up the field of view into nine planes to enable 3D tracking of the rheological probe^25^ and therefore, is not required for LSFM imaging and can be easily removed by means of a slider. Additionally, the multiplane gratings used were optimised for a particular wavelength (543 nm) making them unsuitable for multicolour imaging. OptoRheo is fitted with a stage-top incubator that regulates temperature, CO_2_ and humidity around the sample, allowing long time-course experiments spanning over hours (Fig 2C and Supplementary Videos VS1 and VS2) and days.

### Microrheology of gels and suspension cultures

The viscoelastic properties of biomaterials can be extracted non-invasively using particle tracking microrheology as developed by this group and others^24,25,31^. This involves a statistical analysis of the residual Brownian motion of micron-sized spherical probes, whose temporal behaviour can be described by means of a Generalised Langevin equation^31^. For this purpose, polystyrene microsphere probes were seeded in the cell culture samples under sterile conditions (Fig 3A). For hydrogel-based cell culture systems, the diameter of the microsphere probes (6 μm) was selected so that the Brownian motion of the probes were constrained by the hydrogel polymer network. A small field of view (typically 14 μm x 14 μm) was recorded around the microsphere probe (Fig 3B) to track the trajectory of each probe (Fig 3C) at a relatively high frame rate (~ 300 Hz - 5 kHz) to achieve broadband microrheology. This was done while switching the illumination to transmission mode using a LED source to avoid introducing fluorescence bleaching-related errors in particle tracking (Fig 1). A second cylindrical lens (CY2 in Fig 1) was placed in the LED light path to compensate for the presence of the light sheet forming cylindrical lens (CY1 in Fig 1) and produce uniform illumination. An analysis of the mean squared displacement (MSD) of the confined microspheres (Fig 3D) gives the elastic (*G*’(ω)) and viscous (*G*”(ω)) moduli of the surrounding gel (Fig 3E), see methods section for further details.

**Figure 3:**
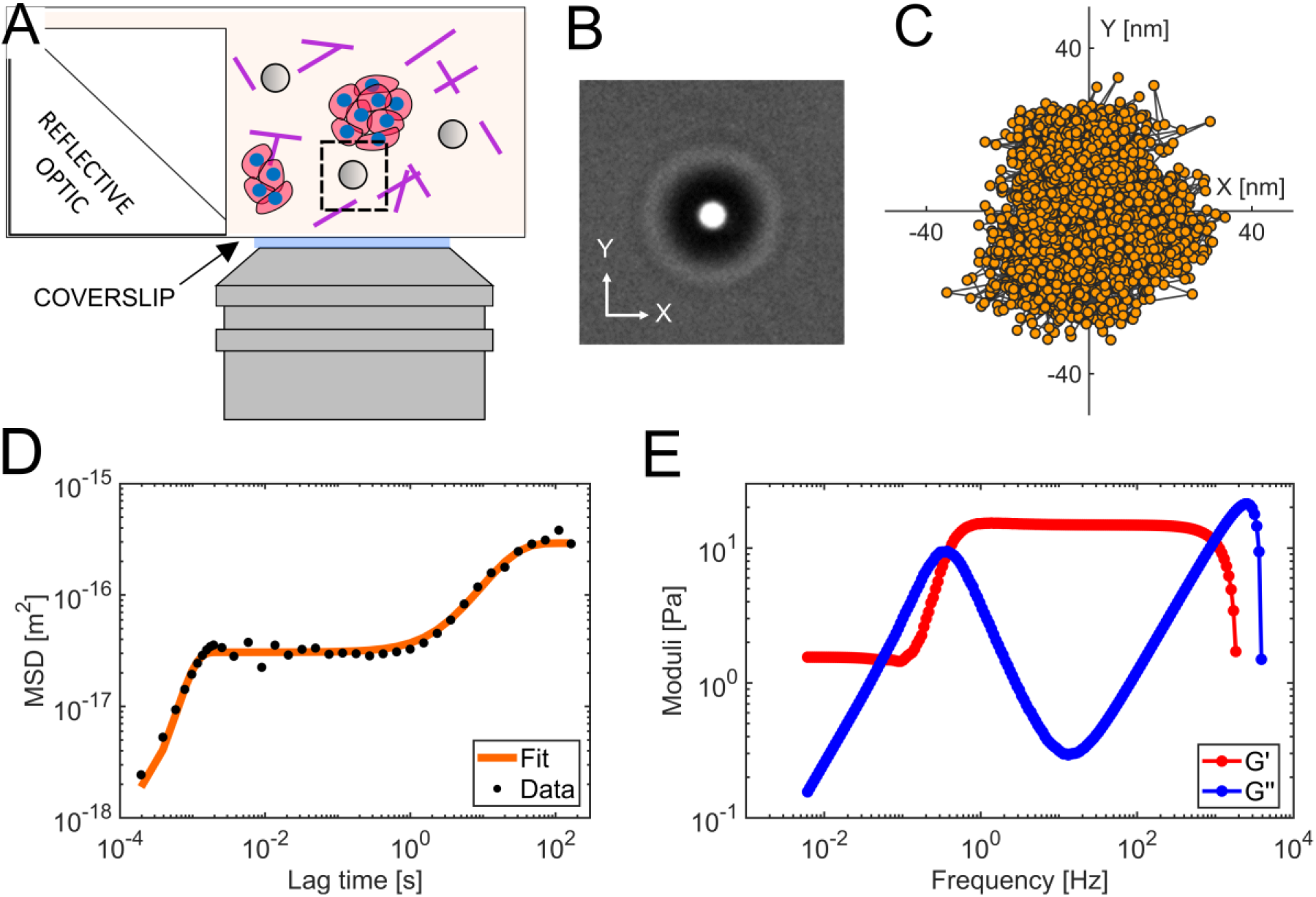
Passive microrheology without optical trapping. A. A schematic representation of passive microrheology measurements. B. Microsphere probes (6 μm diameter) were seeded in the hydrogel and small regions of interest (typically 14 μm x 14 μm as shown) around them were imaged at a high frame rate (5 kHz in this example). C. Particle trajectory in a 2D plane with individual positions depicted in orange (down sampled here for clarity from a total of 1.5 x 10^6^ frames). D-E. The mean squared displacement (MSD) provides a measure of gel compliance, which is then used to compute the elastic (*G*’(ω)) and viscous (*G*”(ω)) moduli of the gel over a wide range of frequencies as shown in panel E (for more details on the analysis see Methods).

In liquid media and suspension cultures, microsphere probes sediment to the coverslip over time with a rate dependent on the gravitational force, the buoyancy of the microsphere and the viscosity of the medium^32^. In aqueous liquids, this sedimentation rate is about 20 μm/s for microspheres with a 3 μm radius, thus preventing long-term (i.e. tens of minutes) tracking of the probe trajectories, which are needed for broadband microrheology calculations.

Therefore, when working with liquids we used an optical trap (Fig 4A) to hold the probe in the field of view and at the required location relative to the cell/s of interest during the measurement time. The trapping force acting on the microsphere was kept very low (<10^-6^N/m) by controlling the laser power, to maximise the amplitude of the residual Brownian motion, increasing the sensitivity of the microrheology measurement and the frequency range. The confined Brownian motion of the microsphere could then be recorded at ~300 Hz in 3D by inserting a removable pair of quadratic gratings (QG) in the detection path before the sCMOS camera to achieve multiplane detection of the particle position^25,33^ (Fig 1 and 4B). In particular, the quadratic grating pair focus light from nine object-planes as an array onto the camera sensor for instantaneous 3D imaging without the need of any mechanical moving parts^25,33^ (Fig 4B). This method allows tracking of the microsphere motion in all three dimensions simultaneously (Fig 4C), revealing spatial variation of the sample’s viscoelastic properties. However, the smallest variance in particle position we could reliably measure with the present configuration was 30 nm in z, compared to 15 nm in x and y. Therefore, tracking in z was unreliable for our gel samples where the motion is typically less than 50 nm (see Fig 3C).

**Figure 4:**
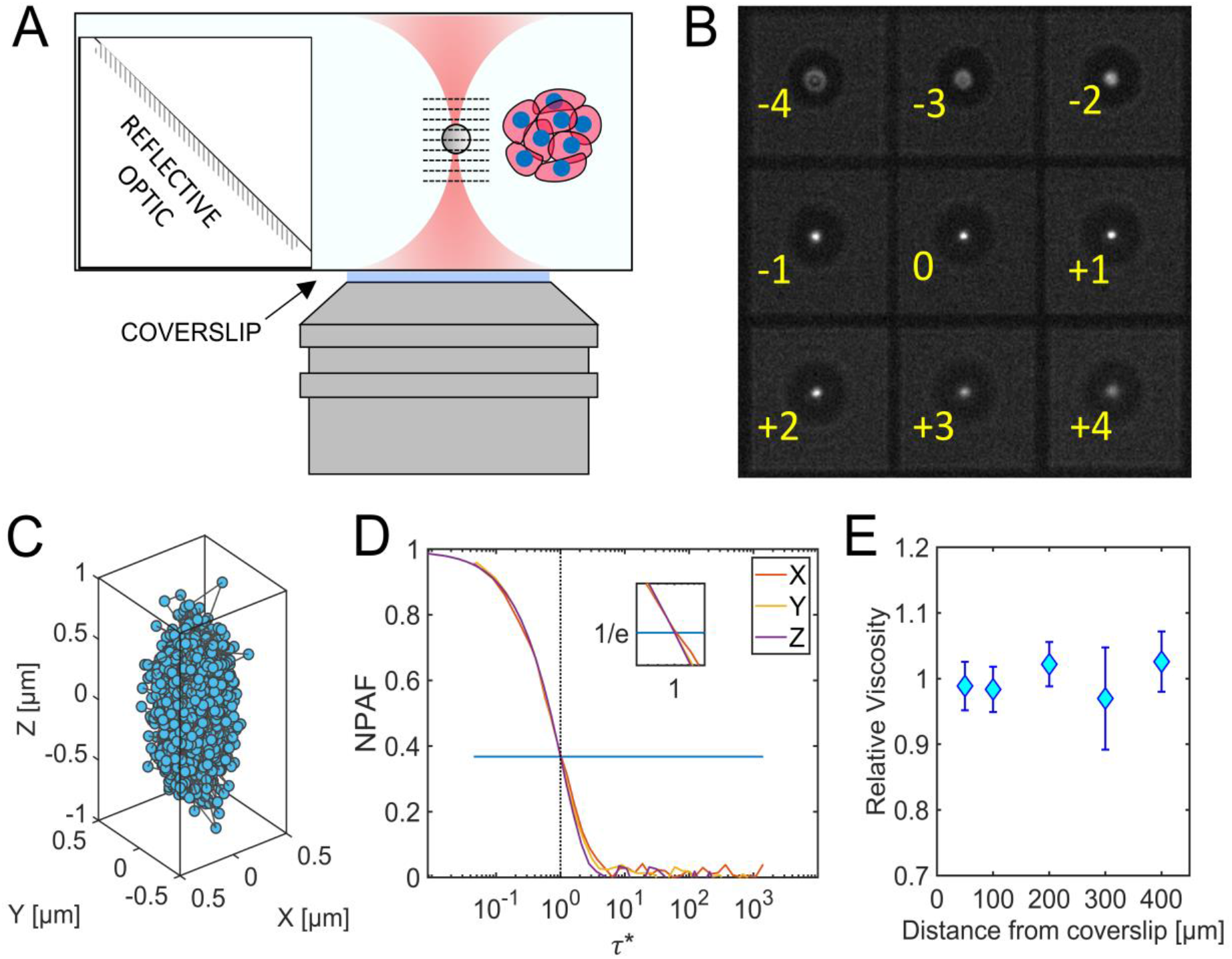
Microrheology with an optical trap and multiplane detection. (3D particle tracking mode) in an aqueous solution. A. An optically trapped microsphere is imaged in 9 planes simultaneously (planes represented by dashed lines). B. Captured image of nine separate Z planes (Δz = 0.79 μm). The planes (labelled from number −4 to +4) are simultaneously recorded at the camera sensor to extract the 3D trajectory. C. The resulting 3D trajectory of optically trapped microsphere in water with individual positions depicted as blue circles (down sampled for clarity from a total of 2 x 10^5^ frames). D. The normalised position autocorrelation function (NPAF) versus a dimensionless time, *τ*\*. E. The mean ± standard deviation of the relative viscosity measured at each position over a range of depths.

We validated our microrheology measurements from the 3D particle tracking mode of OptoRheo by using water, a well characterised Newtonian fluid. We extended our previous work^25^ by measuring the 3D trajectories of microsphere probes from 50 to 400 μm away from the coverslip, without the use of any aberration correction, thus enabling microrheology measurements at the same sample depths as our light-sheet imaging experiments. It is important to note that most studies employing optical trapping report measurements taken at <100 μm from the coverslip^34^. The use of water immersion and a correction collar allowed us to achieve trap stiffness *k* values of *k_x_*= 3.2 x 10^-7^± 0.3 x 10^-7^ N/m along the x axis, *k_y_* = 3.2 x 10^-7^ ± 0.5 x 10^-7^ N/m along the y axis and *k_z_* = 6.7 x 10^-8^ ± 1.2 x 10^--8^ N/m along the z axis (mean ± standard deviation) over this large range of distances from the coverslip.

Relative viscosity (ratio of viscosity of an aqueous solution to the viscosity of water at the same temperature) could be evaluated in 3D by analysing the normalised position autocorrelation function (NPAF) for x, y and z at depths ranging from 50 μm to 400 μm from the coverslip. In particular, the relative viscosity can be read “*at a glance”* from the abscissa of the NPAF intercept e^-1^, when the NPAF is plotted versus a dimensionless lag-time *τ*\* = *kτ*/(6*παη_s_*); where *k* is the trap stiffness, *τ* is the lag-time (or time interval), *α* is the probe radius, and *η_s_* is the Newtonian viscosity expected for the pure solvent^35^ (Fig 4D). Figure 4E shows the mean x, y and z relative viscosity ± standard deviation, performed at different depths (see Methods). Over the range of recorded measurements, the measured mean relative viscosity (over x, y and z) remained stable with depth (i.e., 1 ± 0.05).

### Monitoring ECM stiffness in hydrogel-encapsulated 3D cell culture

In order to test the ability of OptoRheo to evaluate biomechanical properties of the ECM in real time, clusters of human-derived MCF-7 cancer cells expressing the tdTomato fluorescent protein were grown in a hydrogel-encapsulated cell culture matrix^12,36^. Echoing what is known from patients, changes in the ECM stiffness around these cells have been correlated with cancer progression and metastasis, and have been shown to alter drug resistance^37^,^38^. Complementary multi-colour 3D LSFM imaging allowed cells and labelled matrix components (i.e. collagen I) from the same locations to be captured in separate colour channels. These volume images could be overlaid and combined with microrheology measurements to map the changing biomechanical properties to the changing morphology of the sample, at the microscale. This approach was used to compare four different complex systems made of hydrogels with and without (i) collagen and (ii) cells.

The curves of the elastic (*G*’(*ω*)) and the viscous (*G*”(*ω*)) moduli of all the hydrogel preparations show a pattern characteristic of viscoelastic polymer gels^39^ with a rubbery elastic plateau 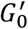 (where *G*’(*ω*) >*G*”(*ω*)) at low frequencies transitioning to a high frequency glassy state (Figs 3E and 5A). Environmental control on OptoRheo allowed the gel samples to be kept under physiological conditions over multiple days, making it possible to re-visit selected microsphere probes multiple times over the duration of the experiment (three days) to follow the changing ECM properties over time. This was performed in triplicate.

**Figure 5:**
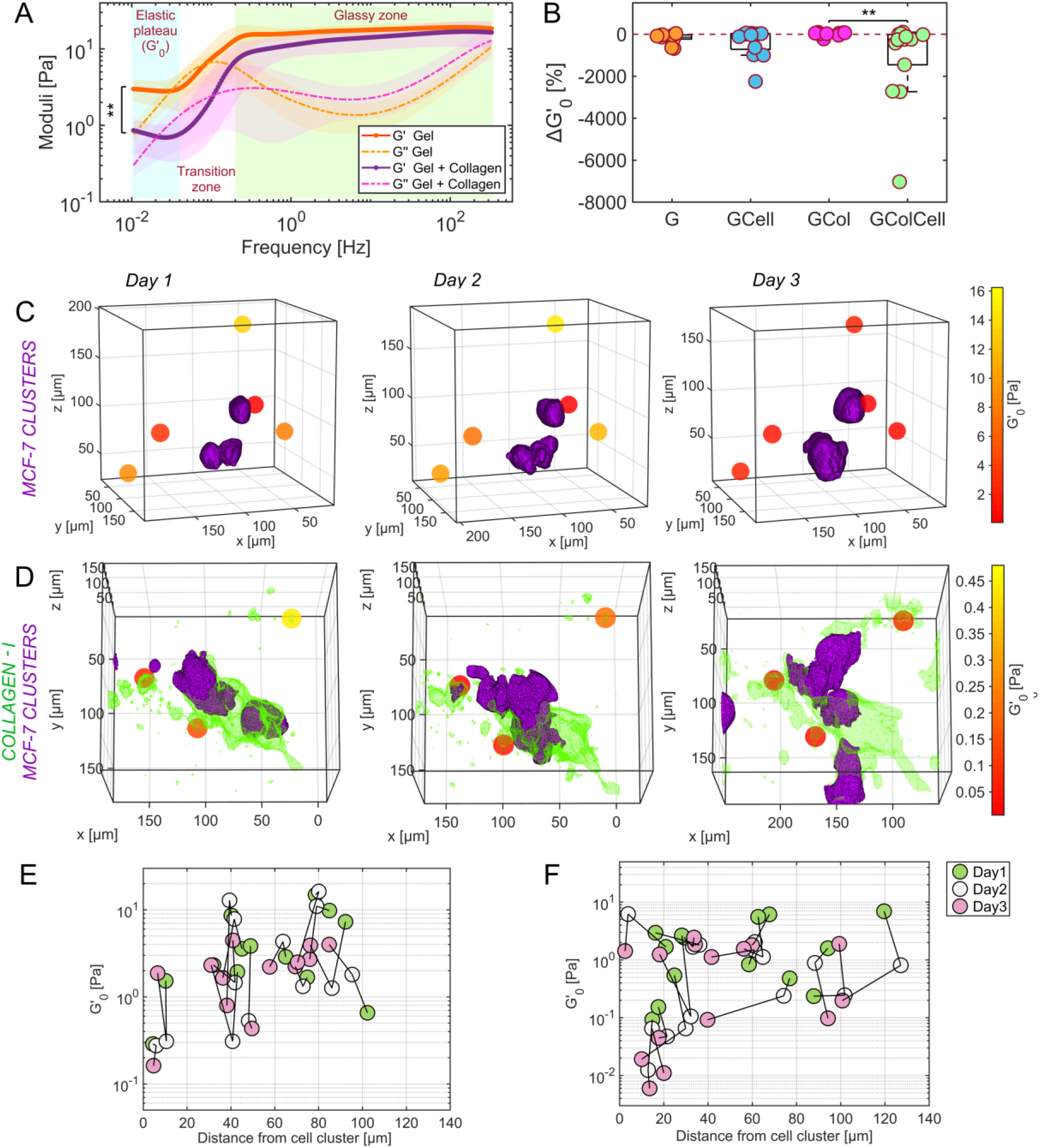
Local stiffness measured in live 3D cell cultures with different compositions. A. Complex moduli of plain gel in orange and gel supplemented with collagen in purple at day 1 (averaged with 95% confidence intervals shown as shaded regions). B. Proportional change in the height of the low frequency elastic plateau 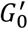 at individual bead probes over three days of observation in plain gel (G), gel with collagen (GCol), gel seeded with cells (GCell) and gel with collagen and cells (GColCell). The dashed line represents no change, negative values indicate more compliant gels. C-D. Biomechanical maps produced by OptoRheo of MCF-7 clusters expressing tdTomato (shown in purple) encapsulated in hydrogels and D. MCF-7 clusters from the same cell line in hydrogel supplemented with collagen I labelled with Cy5 (shown in green) monitored over three days. Spheres depict microsphere probes (not to scale) assigned a colour to reflect the local stiffness 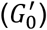. E.& F. Spatiotemporal changes in 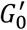 values with relative distance from the edge of the cell clusters in gel in the absence (E) and presence (F) of collagen over three days.

In the absence of cells, the height of the elastic plateau - 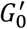, is greater for plain gels (n = 11) as compared to gels supplemented with collagen I (n = 11) (p = 0.0014, Kruskal-Wallis test; Fig 5A) at the start of the experiments (day 1), indicating stiffer gels and may point to differences in the cross-linked network^40^. The high frequency glassy response which can be attributed to local monomer relaxation^41^ is similar for the gels (Fig 5A) which have the same polymer hydrogel base at the same concentration. We therefore focus our analysis on the 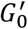 values.

The presence of cells brings about complex changes in the gels which become more apparent over time and when taking into account the proximity to cell clusters (Fig 5 B-F). Over time, as cell clusters proliferated and changed their relative distance from the probes, this information could be extracted from LSFM images. Our data show that the presence of collagen makes the gels more compliant (p = 0.0008 Generalised Mixed Effects Model). There was a significant difference in the proportional change in 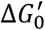 (between day 3 and day 1) in the presence and absence of cells in gels supplemented with collagen (p = 0.007, Kruskal-Wallis test; Fig 5B). Moreover, the 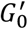 values change with distance from the nearest cell cluster both in the presence and absence of collagen. In the presence of collagen, the gel was most compliant within 50 μm from the edge of the nearest cell cluster. This region near the cell clusters was much more heterogeneous with measurements spanning over three orders of magnitude as compared to farther away (two orders of magnitude at regions > 50 μm) (Fig 5F) hinting at a gel remodelling front.

### Mapping relative viscosity local to spheroids in suspension culture

The second scenario tested as a proof-of-concept, was to acquire 3D images and microrheology measurements near spheroids in suspension culture. Spheroids were grown from the same MCF-7 cancer cell line as in the peptide hydrogel cultures and were used two days after seeding at a size of ~1 mm in diameter. As both 3D imaging and microrheology on OptoRheo do not involve moving the sample, these spheroids could be maintained in liquid media without the need to immobilise them in agarose or any other hydrogel matrix – an advantage over most conventional LSFM instruments. Volume images near the edges of spheroids were acquired by tiling multiple overlapping imaging volumes of 200 μm x 200 μm x 200 μm (between 150 μm – 350 μm from the coverslip) (Fig 6A). Once the images were acquired, the instrument was switched from LSFM modality to 3D particle tracking mode by sliding the quadratic gratings into the optical path and with illumination in transmission (QG in Fig 1). The optical trap enabled microsphere probes to be individually trapped and positioned in 3D with the XY stage and the piezoelectric objective scanner to make measurements at selected locations near the edge of the spheroids (Fig 6A inset).

**Figure 6:**
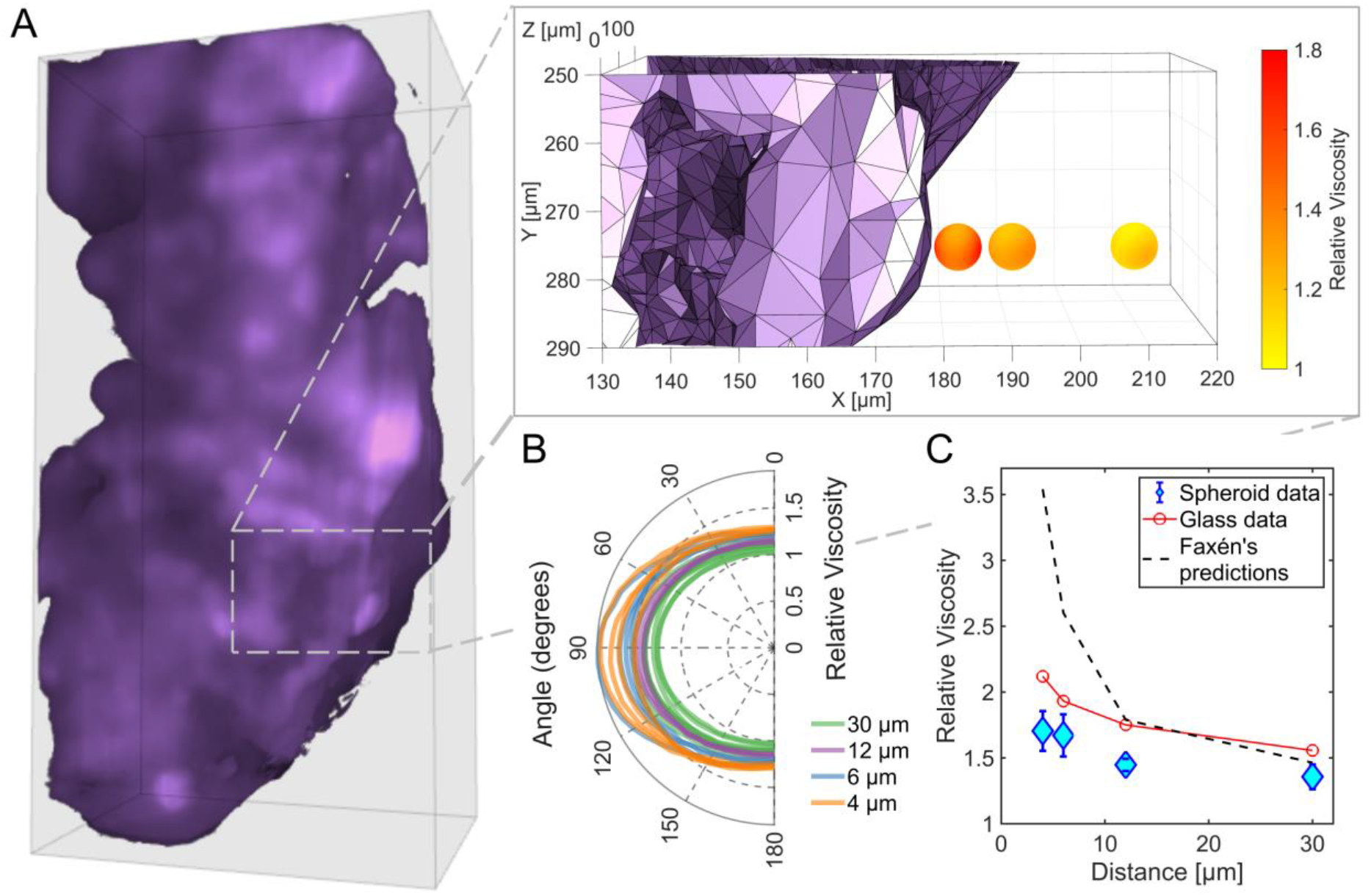
Viscosity near spheroid. A. 3D rendering of a section of a spheroid of MCF-7 cells expressing td-Tomato (relative dimensions: 200 μm x 400 μm x 100 μm (150-250 μm from the coverslip)). The inset shows one of the areas where viscosity (relative to the solvent) measurements were acquired at incremental distances from the spheroid surface – three measurements (4 μm, 12 μm and 30 μm from the spheroid surface) are depicted as spheres (not to scale). The colour gradient for each sphere represents the relative viscosity sampled by angle at each measurement position. A fourth position (6 μm) has not been shown to aid visualisation. B. Relative viscosity measurements in a plane perpendicular to the spheroid surface, passing through the highest and lowest measured viscosity at the probe position. C. Mean ± standard deviation of relative viscosity measurements of the nutrient media at each position for a direction perpendicular to the spheroid surface (blue) and glass (red), showing increased apparent viscosity at positions closer to the surface but lower than values predicted from Faxén’s law (black dashed line).

Our measurements show an apparent increase in relative viscosity with decreasing distance (4 μm, n = 4; 6 μm, n = 4; 12 μm, n = 3 and 30 μm, n = 7) between the centre of the microsphere and the surface of the spheroid (n = 2 spheroids) (Fig 6A, 6B and 6C). Viscosities could be extracted in 3D by resampling the recorded 3D trajectories along any desired axis, calculating the MSD, and then using Fick’s Law (see methods). Such angle-by-angle analysis reveals higher relative viscosity values perpendicular to the surface of the spheroid as compared to parallel to the surface, with the anisotropy increasing as the probe approaches the surface (Fig 6B). This trend is in agreement with predictions from Faxén’s law which describes the increased hydrodynamic drag experienced by objects near solid surfaces and manifests itself as an increase in apparent viscosity, albeit our measured values are lower than the predictions^42^ (dashed black line in Fig 6C). The lower values could potentially be attributed to the spheroid surface being irregular and not completely inelastic. Additionally, the presence of salts in the nutrient medium may screen charge-charge interactions between the microsphere probes and the cell surface, reducing the force required to move the probe closer to the spheroid surface^43^ as corroborated by control measurements at corresponding distances from the inert glass coverslip in the same medium without the presence of the spheroids (red line in Fig 6C).

## Discussion

To understand how cells interact with and remodel their surrounding matrix, it is crucial not only to visualise cell clusters in 3D, but also to map these images to the changing micro-mechanical properties of the matrix local to and distant from the cells. In this study, we have introduced an integrated instrument – OptoRheo, that combines light sheet fluorescence microscopy (LSFM) with non-invasive microrheology to enable a more complete understanding of cell-matrix interactions. The new LSFM configuration presented here is straightforward to implement and does not require the use of bespoke, expensive optics. This reflected configuration allows for samples to be prepared and mounted as on any commercial inverted microscope, using off-the-self sample chambers and a stage-top incubator to control temperature, humidity and CO_2_ throughout experiments, enabling delicate hydrogel-based cell culture samples to be studied over multiple days. With the ability to optically trap microrheological probes when required, we have demonstrated the capability of the instrument to study aqueous (suspension culture) as well as soft solid (hydrogel) environments. This modular functionality with a gratings-based approach allows OptoRheo to transition from 2D to 3D particle tracking without difficulty. We provide experimental evidence of this approach by following changes in matrix viscoelasticity in 2D in hydrogel-encapsulated cell cultures over three days and viscosity in 3D near spheroids in suspension. Our localised rheological measurements reveal heterogeneities at the microscale in hydrogel-encapsulated cell cultures.

OptoRheo provides broadband microrheology measurements, covering 5 - 6 decades of frequency in our data. These broadband measurements provide valuable insight into the frequency-dependent mechanical properties of biological materials. As can be seen in Figure 5, OptoRheo is sensitive to mechanical changes arising from changes in matrix composition and from cell-driven re-organisation of the local environment, detectable over a broad range of frequencies. This is in contrast to Brillouin scattering – another emerging technique that is being used to quantify the mechanical properties of biological systems^44^, but which is limited to a narrow range of frequencies in the gigahertz regime. It is possible that the narrow frequency range of Brillouin scattering measurements will miss some of the mechanical changes we report here. Furthermore, compared to previously published methods for optical trapping based stiffness measurements^45,46^, OptoRheo can characterise samples at depths of hundreds of microns from the coverslip making it particularly suited to cell cultures grown in 3D. OptoRheo has the additional benefit that the mechanical characterisation is paired with multi-channel 3D fluorescence light sheet imaging allowing the changing morphology of the cells to be monitored at the same time.

Extending the rheological measurements to 3D and sampling viscosity in 360 °, as shown in our experiments with spheroids in suspension (Fig 6B), increases the capability of our measurements to extract heterogeneities, not just between probe positions, but for different directions at a single probe position. Currently, this 3D approach is restricted to computing viscosity in liquids due to limited spatial sensitivity when tracking probe position in z (Methods). This is not an issue when the extent of the Brownian motion of the probe is large, such as the ~1 μm (Fig 4B) observed in a weak optical trap in suspension culture, but is of concern when motion is very small (≤ 50 nm) such as that observed in stiff gels (Fig 3B). In future, the sensitivity and measurement range of the ‘sharpness’ metric used for particle tracking in z could be tuned by changing the plane spacing selected with the multiplane grating pair, the probe size, the illumination levels and the signal to noise ratio of the images^25^. Efforts to extend these 3D analyses to gels in the future would be highly valuable as biophysical properties in the ECM are likely to vary in 3D, as apparent in the images of labelled collagen present in the gel samples (Fig 5B).

The outputs from these proof-of-concept experiments are very data-rich. Different locations of interest in the sample can be programmed to be revisited multiple times over a multi-day experiment. As such, OptoRheo enables researchers to track numerous variables over time, so that the relative cell and probe position, and cell behaviour (change in shape or size, migration, apoptosis) can be related to viscous and elastic components of the matrix biophysical properties. In future, it would be relatively straight forward to include additional probes into the sample (either genetically engineered reporters in the cells or sensors embedded in the gel/ matrix) to track the coordinated impact of chemical, biological and mechanical cues.

As highlighted in recent publications^4,47^, the control of cell behaviour by mechanical forces exerted through the ECM remains poorly understood, even as researchers take advantage of ECM control to create more complex, physiologically relevant models of development and disease, that better represent the *in vivo* micro-environment. The ability to integrate read-outs of cell behaviour with the microrheology of pericellular and distant matrix will be critical in further improving these models and using them to uncover the mechanistic basis of the phenomena they imitate. In addition, for development of therapeutics, there remains a significant gap in our understanding of the environments that drugs and delivery vehicles encounter in the body. The simultaneous observation of material transport together with rheological measurements will enable us to build detailed structure-function relations of drug delivery pathways, which in turn, will enable more efficient screening of candidate therapeutics and better predictive models of *in vivo* activity and efficacy.

## Online Methods

### Light sheet fluorescence microscopy (LSFM)

OptoRheo uses a reflected light sheet configuration where light sheet illumination is introduced into the imaging plane of an inverted microscope (Olympus IX-73) using a right-angle optic (90:10 RT beam splitter cube, 10 mm; Thorlabs Inc.). Z-scanning was achieved by moving the light sheet through the sample using a galvanometer scanning mirror (dynAXIS 3S; SCANLAB Gmbh) whilst simultaneously moving the objective lens (LUMFLN60XW 60x 1.1 NA 1.5 mm WD; Olympus) via a motorised piezo objective scanner (P-725.4CD; Physik Instrumente Ltd.) to keep the light sheet in focus. Lateral positioning was achieved using a XY microscope stage (MS-2000, ASI) and a zoom-mount attached to the cylindrical lens forming the light sheet. The fluorescence image was detected using an sCMOS camera (Hamamatsu ORCA Flash 4.0 V2). Environmental control was achieved using an Okolab stage-top incubator (H301-K-FRAME) supplied with pre-mixed CO_2_ gas.

Multi-colour fluorescence imaging was made possible by using three lasers separately to form the light sheet; 473 nm (SLIM-473; Oxxius), 532 nm (BWN-532-2OE; B&W Tek) and 640 nm (OBIS; Coherent). These laser lines were coupled to each other in the illumination beam path using dichroic mirrors. A cylindrical lens (f = 50 mm, Thorlabs) was used to generate the light sheet with a beam waist of 2.6 μm for λ_ex_ 473 nm, 2.4 μm for λ_ex_ 532 nm and 3.3 μm for λ_ex_ 640 nm. The light sheet was aligned and characterised by imaging it in reflection using two beam splitters (90:10 RT beam splitter cube, 5 mm; Thorlabs Inc.) in tandem. The relationship between voltage applied to the galvanometer mirror and position of the light sheet was characterised using this double beam splitter set up and keeping the detection objective stationary while scanning the light sheet in z. The slope of the linear fit to the measured position of the light sheet against the voltage applied gave the pixel to voltage step size for synchronised movement.

When imaging samples, an autofocus step is first performed to ensure the illumination and detection optics, primarily the objective, are aligned. This involves recording a stack of images while keeping the imaging objective stationary and scanning the light sheet with a sub-beam-waist step size. The frame with the highest mean intensity value denotes where the light sheet waist coincides with the imaging plane and so the position of the light sheet for this frame is synchronised with the height of the objective.

Standard off-the-shelf beam splitter cubes have a blunt edge that make the bottom 150 μm unsuitable for reflecting the light sheet illumination. These regions can be illuminated by tilting the light sheet at BS1 (Fig 1) or by using a bespoke cube with a sharp edge.

The mechanical components of the OptoRheo including the light sheet parts were controlled in LabVIEW (2018, 64bit; National Instruments Inc.). Image volumes were saved as ‘.tiff’ files. Automation of the laser lines through camera-controlled remote triggering and a motorised filter turret enabled overnight time lapse imaging.

### Image processing

Contrast adjustment and background subtraction was performed on image volumes in ImageJ/ Fiji^48^ and volume tile stitching was performed using the BigStitcher^49^ plugin for Fiji. 3D rendering for Figure 6A was done in FluoRender (v 2.26.3)^50^ and for Figure 2B was done in Imaris (10.0.0, Oxford Instruments).

To calculate the distance between microspheres and the nearest cell clusters, the centre positions of the microspheres in image coordinate space were extracted from the LSFM images. Although the microspheres (Polybead^®^ Microspheres 6.00 μm; PolySciences) were not fluorescently labelled, they are identifiable in the 3D LSFM images due to light scattering. Mesh renderings of the corresponding cell clusters were exported from FluoRender and these meshes along with positions of the microspheres from the same image volume were used as inputs in the point2trimesh.m^51^ code in MATLAB which computes the shortest distance between a given point and the outer edge of a triangular mesh.

Figures 5C and 5D and the inset within 6A were prepared in MATLAB using mesh renderings generated in FluoRender overlaid with rendered spheres to depict the microsphere probes with a colour gradient to show the low frequency plateau in the elastic modulus 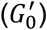 (Fig 5C and 5D) or viscosity (Fig 6A inset) at each probe.

### Optical Tweezers

The beam path from a continuous wave 1064 nm 5 W DPSS laser (Opus, Laser Quantum) was directed into the inverted microscope body and focused in the image plane using the same objective lens used for imaging in the LSFM set up. This objective lens was also used to image a small region of interest (14 μm x 14 μm), required for high frame rate imaging, around the trapped polystyrene microspheres in wide-field with illumination in transmission for fast (300 Hz for multiplane 3D rheology) tracking of thermal fluctuations.

### Multiplane detection

3D imaging of the microrheology probes was made possible by multiplane detection similar to the OpTIMuM instrument^25^ and its predecessors^33,52^. Here, a multiplane grating pair was formed using two quadratically distorted diffraction gratings etched into a quartz substrate (bespoke production by Photronics UK Ltd). A single grating generates three sub-images, corresponding to the m = 0, ± 1 diffraction orders while two gratings with orthogonal etch patterns, can generate nine different sub-images, each corresponding to a different image depth which can be captured simultaneously on a single camera sensor (Hamamatsu ORCA Flash 4.0 V2) (Fig 4A). A 4f image relay system consisting of two 300 mm lenses was set up in the detection path between the camera and the inverted microscope body to enable the multiplane grating pair to be placed in the telecentric position. This set up ensured a consistent level of magnification in each of the imaging focal planes. The grating is on a slider and easily removable allowing the user to switch between standard full field of view imaging and multiplane imaging of a small region of interest with no adverse side effects. In our system we have used a relay and grating combination that gives plane separation of Δz = 0.79 μm with the nine images spanning 7.11 μm, designed to show the extent and position of our 6 μm diameter probe. Grating combinations can be chosen to suit the diameter of the probe such that the total span in z covers the extent of the trajectory of the probe with the minimum plane separation for optimal resolution^25^.

### Microrheology

Particle tracking microrheology was performed using polystyrene microspheres as probes (Polybead^®^ Microspheres 6.00 μm diameter; PolySciences). In hydrogel cultures, the microspheres were encapsulated during the gelation process at a final density of 3 x 10^5^ particles/ mL. In suspension cultures, the microspheres were added to a final dilution of 1:200,000 from concentrate product, the probes were individually optically trapped using ~4 mW of laser power (at the sample) and moved to a position of interest. The Brownian motion of the microspheres, was recorded over 300,000 frames at ~300 frames per second for experiments in Figures 4 and 5, and for 1.5 x 10^6^ frames at 5 kHz for Figure 3 and the measurements depicted in the time lapse videos (VS1 and VS2) using OptoRheo with illumination in transmission from an LED light source (Fig 1). Videos of the microsphere probes were acquired using Micro-Manager (version 1.4)^53^ and Micro-Manager (version 2.0) for Figure 3 and the measurements corresponding to the time lapse videos.

The time-dependent trajectories of the microspheres were extracted from these videos in MATLAB (2019b; MathWorks, Nattick, MA). For 2D trajectories along the image plane a centre-of-mass detection method following Otsu’s method of multiple thresholding (with two levels) was used. Out-of-plane Z motion of the probe was tracked by computing a ‘Sharpness’ metric as detailed in our OpTIMuM publication^25^. Particle tracking with these methods gives us a minimum sensitivity of ~ 15 nm (FWHM) in the xy plane and ~ 30 nm (FWHM) in z for a particle with diameter of ~ 6 μm, using a 60x objective and a plane spacing of ~ Δz = 0.79 μm^25^. A calibration step is performed for each microsphere before taking a measurement by translating a lens (L4 in Fig 1) in the beam expander in the optical path as described previously^25^.

In the case of hydrogels, the Brownian motion of the microsphere confined within the gel was recorded in 2D without the use of the optical trap or multiplane imaging. For these data, an analysis of the mean squared displacement (MSD) gave the storage (elastic) and loss (viscous) moduli of the gel. To acquire the viscoelastic measurements for each probe, first each experimentally acquired trajectory was detrended to remove long-term drift and a filtering step was performed to remove instrument noise. For the noise filtering, a Fourier transform of each trajectory was used to identify sharp noise peaks characterised by a single frequency width and using an amplitude threshold of 2 x 10^-10^ m. These noisy peaks, attributed to electrical noise from the laboratory, were then removed from the data using a custom multiband filter in MATLAB. The MSD values for each filtered trajectory was then fit with a stretched bi-exponential of the form

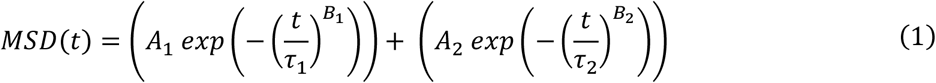

A_1_, A_2_, B_1_, B_2_, τ_1_ and τ_2_ are fitting parameters. This approach mitigates any error generated by the inherently finite nature of the measurements that affects the accuracy to which the MSD is calculated especially at short-time scales (Fig 3). The MSD relates to the gel’s time dependent compliance J(t)^54^ as follows,

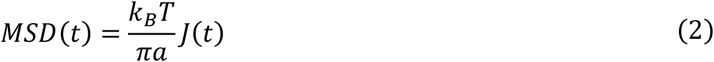

where k_B_ is the Boltzmann’s constant, T is the absolute temperature, and a is the radius of the microsphere. The materials’ complex shear modulus can be computed from the materials’ compliance by means of its Fourier transform 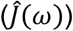

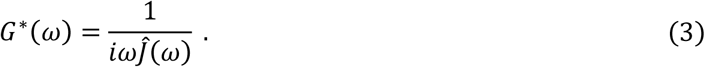

We used a new MATLAB based graphical user interface named π-Rheo (see code availability statement) for evaluating Equations 2 & 3, to compute the Fourier transform of the particles’ MSD and the materials’ complex modulus for passive microrheology measurements. π-Rheo is underpinned by the algorithm introduced in i-RheoFT^55^. The real and imaginary parts of the complex modulus give the elastic (*G*’(*ω*)) and viscous (*G*”(*ω*)) moduli of the gel.

The elastic plateau of the gels 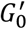 (equivalent to *G*’(*ω*) at low frequencies) can be calculated simply from the particles’ time-independent variance 〈*r*^2^〉 using the formula:

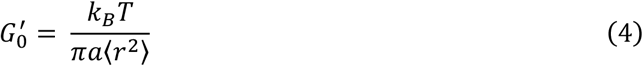

For aqueous solutions, where 3D positions of the probe are tracked, the viscosity may be extracted by fitting an exponential decay against the normalised position autocorrelation function^35^. This method is highly effective for data aligned with the principal axes of the optical trap (see Fig 4D). However, when calculating viscosity along vectors not aligned with these axes using this method, the significant trap anisotropy along the z-axis introduces artefacts as outlined in detail previously^56^. Alternatively, if the material under investigation is purely viscous, then at very early times the MSD of the bead should behave as if the bead is not trapped. Under these conditions, Fick’s Law for unconstrained diffusion can be used to extract viscosity in any arbitrary direction rather than just x, y, z at these early times. Fick’s law for motion in 1D is given by,

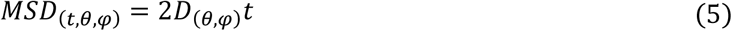

where θ and ψ define the direction being probed and D is the diffusion coefficient for a sphere of radius a in a liquid with viscosity η. From the Stokes-Einstein relation

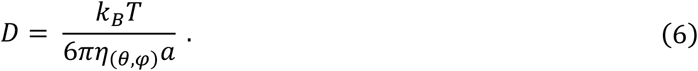

This approach was used to compute viscosity in 3D as shown in Figure 4E and Figure 6A.

### Cell culture

The breast cancer cell lines MCF-7 and MDA-MB-231 expressing tdTomato or eGFP were produced by lentiviral transduction of cells originally obtained under MTA from NCI as part of the NCI-60 panel. These cell lines were maintained in high glucose DMEM (MCF-7 tdTomato; Life Technologies, 21969-035) or phenol red free RPMI (MDA-MB-231 tdTomato and MCF-7 eGFP; Sigma, D5671) with 10 % foetal bovine serum (Life Technologies, 10500-064), 1 % L-glutamine (Life Technologies, 25030-024). To maintain the tdTomato protein expression, the medium was supplemented with Puromycin (Gibco, A11138-03) at 1:1000 every passage (MCF-7 tdTomato) or every 3 weeks at 1:500 (MCF-7 eGFP) or 1:250 (MDA-MB-231 tdTomato). Cells were maintained at 37 °C and 5 % CO_2_ in a humidified atmosphere during cell culture and measurements on the OptoRheo. All cell lines were subjected to monthly mycoplasma testing and none of the lines used in this study tested positive at any point.

### Peptide gel precursor preparation

The precursor and gel preparation method was followed as previously published^57^. A commercially available peptide preparation in powder form was used as the source of the octapeptide gelator (Pepceuticals UK, FEFEFKFK, Phe-Glu-Phe-Glu-Phe-Lys-Phe-Lys). To form the precursor, a mass of 10 mg peptide preparation was dissolved in 800 μL sterile water (Sigma, W3500), using a 3 min vortex step followed by centrifugation (3 min at 1000 rpm) and a 2 hour incubation at 80 °C. After incubation, 0.5 M NaOH (Sigma, S2770) was added incrementally to the gel until optically clear. The gel was vortexed, buffered by addition of 100 μL 10x PBS (Gibco, 70011), and incubated at 80 °C overnight. The resulting precursor could be stored at 4 °C until required.

### Peptide gel formation with collagen I supplementation

Prior to peptide gel formation, the precursor was heated at 80 °C until liquid to ensure homogeneity, before transferring to a 37 °C water bath. Cy5 labelled (in-house preparation, see method below) rat tail Collagen I was neutralised directly before use with 1 M NaOH according to manufacturer instructions, and diluted with sterile water and 10x PBS to a concentration of 1 mg/ mL while keeping on ice at all times to prevent polymerisation. Peptide gel formation was then induced by pH neutralisation on addition of cell culture medium (with or without cell suspension) to the gel precursor. A final volume of 1.25 mL was obtained from a preparation by adding 125 μL of cell suspension and 125 μL Cy5 collagen I to a precursor volume of 1 mL. The end concentration of peptide preparation was 8 mg/ mL and collagen I concentration was 100 μg/ mL. Polystyrene microspheres (Polybead^®^ Microspheres 6.00 μm; PolySciences) were added at final density of approx. 3 x 10^5^particles/ mL. The medium / cell suspension was thoroughly mixed with the precursor and Collagen-I by gentle (reverse) pipetting, before plating at 100 μL per well into a 4 μ-well glass bottom chambered coverslips (IBIDI, 80427) pre-mounted with a beamspliter cube (ThorLabs, BS070). The wells were then flooded with cell culture medium and incubated at 37 °C and 5 % CO_2_ in a humidified atmosphere. Sequential media changes (at least two) over the next 2 hours ensured complete neutralisation and therefore gelation.

For cell encapsulation, the 125 μL volume of cell culture medium was prepared as a cell suspension at 10x the intended final seeding density, to allow for the dilution factor on mixing with the gel precursor. Trypsin-EDTA (0.25%; Life Technologies, 25200056) was used to detach cells from 2D culture at sub-confluence. 1.25 x 10^5^ cells were re-suspended in 125 μL cell culture medium, giving final seeding density in the peptide gel 1 x 10^5^ cells/ mL. For data in Figure 2B, a co-culture of MCF-7 (eGFP) and MDA-MB-231 (tdTomato) were at a final seeding density of 1 x 10^6^ cells/ mL for each cell type. 24 hrs post encapsulation culture medium was replenished, with the addition of HEPES buffer (Life Technologies, 15630-056) at 10 mM final concentration and 0.5 - 1 % penicillin/ streptomycin (Gibco, 15140122).

Prior to casting the gel, the beam splitter cubes were sterilised in absolute ethanol. Cubes were soaked for 1 hour, then left to dry on a paper tissue inside the class 2 safety cabinet. To minimise movement and consequently damage to delicate structure of a hydrogel, the cubes were secured in place with glass coverslips.

### Collagen labelling with Cy5

Rat tail collagen type 1 solution (10 mL; Gibco, A1048301) was mixed with 0.1 M sodium bicarbonate buffer (10 mL, pH 8.5) and 110 μL Cy5 NHS ester solution (10 mg/ mL, DMSO) added. The reaction mixture was stirred at 4 °C overnight. The reaction mixture was purified via the dialysis method at 4 °C to remove the unreacted dye and yield the Cy5 labelled collagen. It was then lyophilised and reconstituted in 20 mM acetic acid buffer.

### Spheroid preparation

Corning 7007 Ultra-low attachment (ULA) 96-well round-bottom plates were used to culture the 3D spheroids. 80 % confluent tdTomato MCF-7 monolayer cells were detached, collected and the cell number determined using an automated cell counter (Biorad TC20). A single-cell suspension was diluted in culture medium and cells seeded at 6000 cells/ well to generate the spheroids (final volume of cell suspension in each well was 100 μL). The plates were then centrifuged at 300 RCF for 5 min and cultured for 3 days until visible spheroid formation.

For experiments on the OptoRheo, spheroids were placed in 4 μ-well glass bottom chambered coverslips (IBIDI, 80427) using a P1000 pipette with the pipette tip cut off at the end. Each spheroid was placed alone in 500 μL of phenol red-free culture media (1:1 DMEM:F12 supplemented with 10% FBS), ~ 5 mm away from the edge of a 10 mm beam splitter cube (ThorLabs, BS070) (Fig S1A) in each well to enable LSFM imaging. Similar to the peptide gel sample preparation protocol, beam splitter cubes were sterilised between uses and secured in place in the chambered coverslips by wedging glass coverslips between the cube and the chamber side wall.

### Time lapse experiment

Time lapse videos were generated of MDA-MB231 cells expressing tdTomato fluorescent protein seeded in peptide gel and supplemented with unlabelled collagen type I (Gibco, A1048301) and bead probes (Polybead^®^ Microspheres 6.00 μm; PolySciences). TrypLE (Gibco, 12604) was used to detach cells from 2D culture at sub-confluence. The gels were prepared in ibidi 4-well chambered coverslips with a beam splitter cube inserted at one end, similar to the gel rheology experiments described above but with final cell seeding density increased to 2 x 10^6^ cells/ mL for video VS1 and 1 x 10^6^ cells/ mL for video VS2, collagen I increased to 150 μg/ mL and bead density increased to 3×10^5^ / mL for samples in both videos. The samples were kept at 37 °C and supplied with humidified 5 % CO_2_ using cell culture incubators prior to imaging and then the Okolab stage-top incubator during the experiment.

Single channel image volumes were acquired at 10 minute time intervals with light sheet illumination at 532 nm using an automated LabVIEW program on the OptoRheo. Acquired image volumes were subjected to 3D deconvolution using the Wiener Filter Preconditioned Landweber (WPL) method in the Parallel Iterative Deconvolution plugin in Fiji. The 4D videos were rendered in Imaris (10.0.0, Oxford Instruments).

The time lapse videos were halted at regular intervals to acquire microrheology measurements within the same field of view by recording the Brownian motion of 6 μm bead probes at 5 kHz for 1.5 x 10^6^ frames. The bead trajectories were analysed as described in the microrheology section above and depicted as spheres (to scale) in the videos by creating volume objects in a separate colour channel with a colour shade corresponding to the local measurement at the time.

## Acknowledgements

The authors acknowledge support via linked EPSRC grants EP/R035067/1, EP/R035563/1, and EP/R035156/1, pilot grant funding from Nottingham Breast Cancer Research Centre, Anne McLaren fellowship funding from the University of Nottingham (JCA) and NC3Rs grants NC/T001259/1 and NC/T001267/1. The MCF-7 eGFP, MCF7-tdTomato and MDA-MB-231 tdTomato cell lines were provided by and with thanks to Prof. Anna Grabowska, University of Nottingham.

## Supplementary Information

### 1. Sample preparation

**Fig S1:**
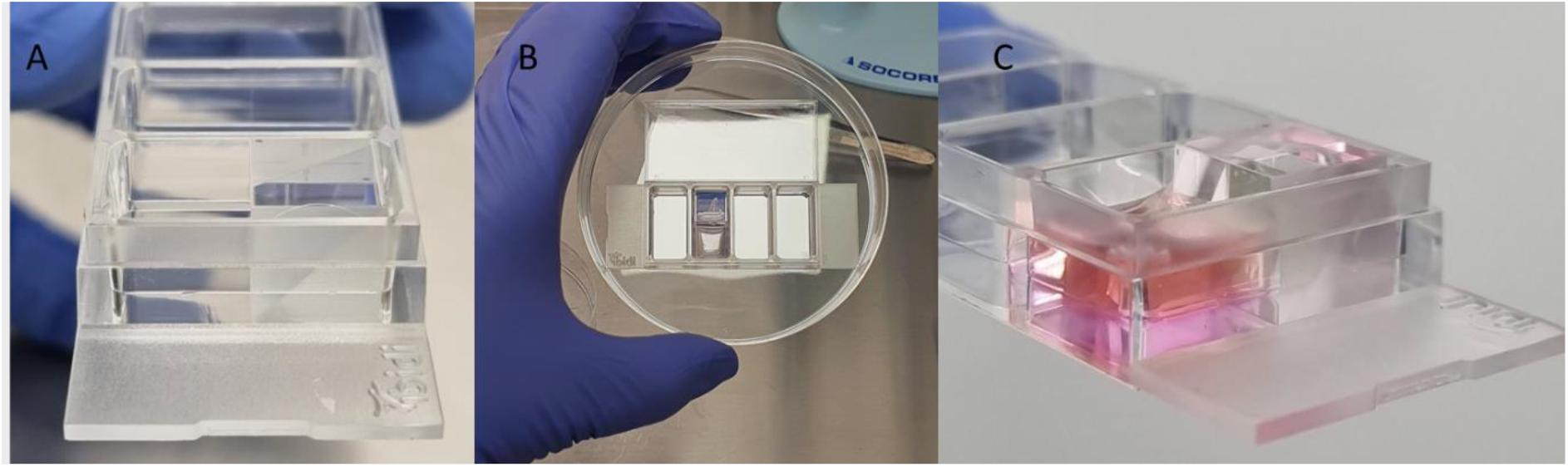
Sample set up: A. Side view of the 4 μ-well chambered coverslip with a 10 mm beam splitter cube inserted with the reflective surface facing the empty half of the chamber. B. Top view of a sample with the gel cast next to the beam splitter cube. C. Side view of the peptide hydrogel topped up with medium next to the beam splitter cube.

### 2. Light sheet properties

**Fig S2:**
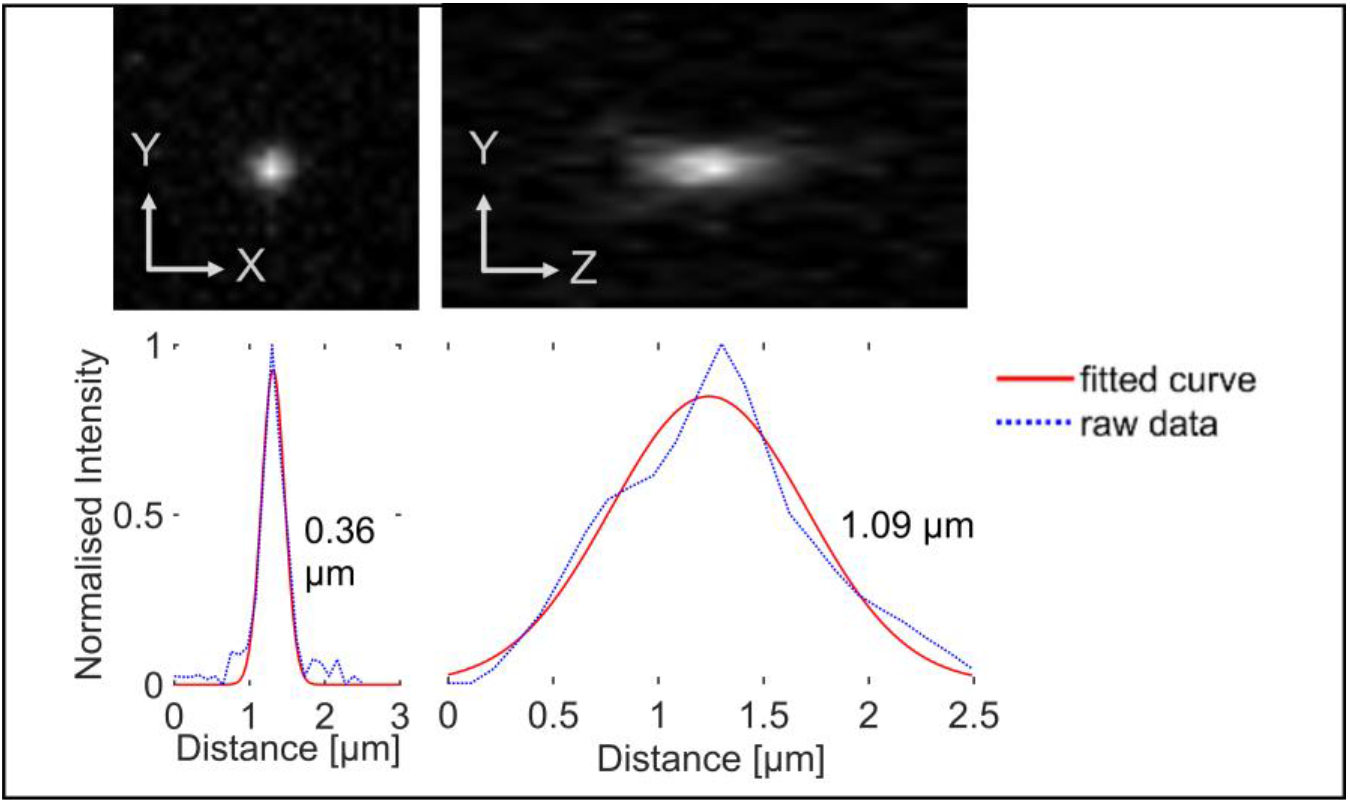
Lateral (left) and axial (right) point spread functions (PSFs) of the system along the XY (0.36 μm FWHM) and YZ (1.09 μm FWHM) planes measured using fluorescent sub-diffraction sized microspheres (diameter = 200 nm, λe× / λem = 532 nm / 580 nm) at ~200 μm from the coverslip.

### Supplementary videos

VS1: **MDA-MB-231 (tdTomato) cells changing morphology** in 3D within a hydrogel matrix supplemented with collagen I (unlabelled). The video was acquired over ~7 hours with a 10 min time interval between frames. Changes in ECM rheology and cell morphology appear related as a more compliant gel at the start of the video (see table S1 below) precedes cell elongation while an increase in stiffness around 6 hours into the experiment corresponds with a retracted cell morphology.

**Table S1:** Microrheology measurements depicted in supplementary video VS1 clockwise from bottom over the time course of the experiment.

| Time | Measurement<br>location 1 ($G'_0$ [Pa]) | Measurement<br>location 2 ($G'_0$ [Pa]) | Measurement<br>location 3 ($G'_0$ [Pa]) |
| --- | --- | --- | --- |
| 0 min | $2.3 \times 10^{-2}$ | $2.0 \times 10^{-2}$ | $1.5 \times 10^{-2}$ |
| 120 min | $1.5 \times 10^{-2}$ | $0.7 \times 10^{-2}$ | $0.4 \times 10^{-2}$ |
| 240 min | $1.2 \times 10^{-2}$ | $2.2 \times 10^{-2}$ | $3.2 \times 10^{-2}$ |
| 360 min | $20.2 \times 10^{-2}$ | $22.9 \times 10^{-2}$ | $24.0 \times 10^{-2}$ |

VS2: **MDA-MB-231 (tdTomato) cells migrating** in 3D within a hydrogel matrix supplemented with collagen I (unlabelled). The video was acquired over 4 hours with a 10 min time interval between frames. Rheology measurements showed a more compliant region (2 x 10^-2^ Pa) near (~ 50 μm) the migratory path depicted as a dark pink sphere as opposed to farther away (6 x 10^-2^ Pa at ~80 μm away) depicted as a bright pink sphere.

## References

1. Discher, D. E., Janmey, P. & Wang, Y. Tissue Cells Feel and Respond to the Stiffness of Their Substrate. Science 310, 1139–1143 (2005).

2. Lo, C. M., Wang, H. B., Dembo, M. & Wang, Y. L. Cell movement is guided by the rigidity of the substrate. Biophys J 79, 144–152 (2000).

3. Pelham, R. J. & Wang, Y. Cell locomotion and focal adhesions are regulated by substrate flexibility. PNAS 94, 13661–13665 (1997).

4. Veenvliet, J. V., Lenne, P.-F., Turner, D. A., Nachman, I. & Trivedi, V. Sculpting with stem cells: how models of embryo development take shape. Development 148, dev192914 (2021).

5. Fernandez, P. & Bausch, A. R. The compaction of gels by cells: a case of collective mechanical activity. Integrative Biology 1, 252–259 (2009).

6. Bloom, R. J., George, J. P., Celedon, A., Sun, S. X. & Wirtz, D. Mapping Local Matrix Remodeling Induced by a Migrating Tumor Cell Using Three-Dimensional Multiple-Particle Tracking. Biophysical Journal 95, 4077–4088 (2008).

7. Lu, P., Weaver, V. M. & Werb, Z. The extracellular matrix: A dynamic niche in cancer progression. Journal of Cell Biology 196, 395–406 (2012).

8. Long, Y., Niu, Y., Liang, K. & Du, Y. Mechanical communication in fibrosis progression. Trends in Cell Biology 32, 70–90 (2022).

9. Meng, H. & Nel, A. E. Use of Nano Engineered Approaches to Overcome the Stromal Barrier in Pancreatic Cancer. Adv Drug Deliv Rev 130, 50–57 (2018).

10. Piersma, B., Hayward, M. K. & Weaver, V. M. Fibrosis and cancer: A strained relationship. Biochim Biophys Acta Rev Cancer 1873, 188356 (2020).

11. Gong, Z. et al. Matching material and cellular timescales maximizes cell spreading on viscoelastic substrates. PNAS 115, E2686–E2695 (2018).

12. Ashworth, J. C. et al. Peptide gels of fully-defined composition and mechanics for probing cell-cell and cell-matrix interactions in vitro. Matrix Biology 85–86, 15–33 (2019).

13. Chaudhuri, O. Viscoelastic hydrogels for 3D cell culture. Biomaterials Science 5, 1480–1490 (2017).

14. Chaudhuri, O., Cooper-White, J., Janmey, P. A., Mooney, D. J. & Shenoy, V. B. Effects of extracellular matrix viscoelasticity on cellular behaviour. Nature 584, 535–546 (2020).

15. Charrier, E. E., Pogoda, K., Wells, R. G. & Janmey, P. A. Control of cell morphology and differentiation by substrates with independently tunable elasticity and viscous dissipation. Nat Commun 9, 449 (2018).

16. Greiss, F., Deligiannaki, M., Jung, C., Gaul, U. & Braun, D. Single-Molecule Imaging in Living Drosophila Embryos with Reflected Light-Sheet Microscopy. Biophysical Journal 110, 939–946 (2016).

17. Beicker, K., O’Brien, E. T., Falvo, M. R. & Superfine, R. Vertical Light Sheet Enhanced Side-View Imaging for AFM Cell Mechanics Studies. Scientific Reports 8, 1504 (2018).

18. Kashekodi, A. B., Meinert, T., Michiels, R. & Rohrbach, A. Miniature scanning light-sheet illumination implemented in a conventional microscope. Biomedical Optics Express 9, 4263 (2018).

19. Gustavsson, A. K., Petrov, P. N., Lee, M. Y., Shechtman, Y. & Moerner, W. E. 3D single-molecule super-resolution microscopy with a tilted light sheet. Nature Communications 9, 123 (2018).

20. Buchmann, B. et al. Mechanical plasticity of collagen directs branch elongation in human mammary gland organoids. Nature Communications 12, 2759 (2021).

21. Hafner, J. et al. Monitoring matrix remodeling in the cellular microenvironment using microrheology for complex cellular systems. Acta Biomaterialia 111, 254–266 (2020).

22. Ciccone, G. et al. What Caging Force Cells Feel in 3D Hydrogels: A Rheological Perspective. Advanced Healthcare Materials 9, 2000517 (2020).

23. Han, Y. L. et al. Cell swelling, softening and invasion in a three-dimensional breast cancer model. Nature Physics 16, 101–108 (2020).

24. Guadayol, Ò. et al. Microrheology reveals microscale viscosity gradients in planktonic systems. PNAS 118, e2011389118 (2021).

25. Matheson, A. B. et al. Optical Tweezers with Integrated Multiplane Microscopy (OpTIMuM): a new tool for 3D microrheology. Scientific Reports 11, 5614 (2021).

26. Huisken, J., Swoger, J., Del Bene, F., Wittbrodt, J. & Stelzer, E. H. K. Optical Sectioning Deep Inside Live Embryos by Selective Plane Illumination Microscopy. Science 305, 1007–1009 (2004).

27. Pitrone, P. G. et al. OpenSPIM: an open-access light-sheet microscopy platform. Nature methods 10, 598–599 (2013).

28. Reynaud, E. G., Krzic, U., Greger, K. & Stelzer, E. H. K. Light sheet-based fluorescence microscopy: more dimensions, more photons, and less photodamage. HFSP journal 2, 266–75 (2008).

29. Hu, Y. S. et al. Light-sheet Bayesian microscopy enables deep-cell super-resolution imaging of heterochromatin in live human embryonic stem cells. Optical Nanoscopy 2, 7 (2013).

30. Gebhardt, J. C. M. et al. Single-molecule imaging of transcription factor binding to DNA in live mammalian cells. Nature Methods 10, 421–426 (2013).

31. Tassieri, M. Microrheology with optical tweezers: peaks & troughs. Current Opinion in Colloid and Interface Science 43, 39–51 (2019).

32. Lee, M. P., Padgett, M. J., Phillips, D., Gibson, G. M. & Tassieri, M. Dynamic stereo microscopy for studying particle sedimentation. Optics Express 22, 4671 (2014).

33. Blanchard, P. M. & Greenaway, A. H. Simultaneous multiplane imaging with a distorted diffraction grating. Appl. Opt. 38, 6692 (1999).

34. Dasgupta, R., Verma, R. S., Ahlawat, S., Chaturvedi, D. & Gupta, P. K. Long-distance axial trapping with Laguerre–Gaussian beams. Appl. Opt., AO 50, 1469–1476 (2011).

35. Tassieri, M. et al. Microrheology with Optical Tweezers: Measuring the relative viscosity of solutions ‘at a glance’. Scientific Reports 5, 8831 (2015).

36. Pal, A. et al. A 3D Heterotypic Breast Cancer Model Demonstrates a Role for Mesenchymal Stem Cells in Driving a Proliferative and Invasive Phenotype. Cancers 12, 2290 (2020).

37. Vasudevan, J., Lim, C. T. & Fernandez, J. G. Cell Migration and Breast Cancer Metastasis in Biomimetic Extracellular Matrices with Independently Tunable Stiffness. Advanced Functional Materials 30, 2005383 (2020).

38. Lovitt, C. J., Shelper, T. B. & Avery, V. M. Doxorubicin resistance in breast cancer cells is mediated by extracellular matrix proteins. BMC Cancer 18, 41 (2018).

39. Shin, M. et al. Rheological criteria for distinguishing self-healing and non-self-healing hydrogels. Polymer 229, 123969 (2021).

40. Abidine, Y. et al. Physical properties of polyacrylamide gels probed by AFM and rheology. EPL 109, 38003 (2015).

41. Cai, P. C. et al. Rheological Characterization and Theoretical Modeling Establish Molecular Design Rules for Tailored Dynamically Associating Polymers. ACS Cent. Sci. 8, 1318–1327 (2022).

42. Leach, J. et al. Comparison of Faxén’s correction for a microsphere translating or rotating near a surface. Physical Review E 79, 026301 (2009).

43. Meza, J. M. H. et al. Particle/wall electroviscous effects at the micron scale: comparison between experiments, analytical and numerical models. J. Phys.: Condens. Matter 34, 094001 (2021).

44. Prevedel, R., Diz-Muñoz, A., Ruocco, G. & Antonacci, G. Brillouin microscopy: an emerging tool for mechanobiology. Nat Methods 16, 969–977 (2019).

45. Rohrbach, A., Tischer, C., Neumayer, D., Florin, E.-L. & Stelzer, E. H. K. Trapping and tracking a local probe with a photonic force microscope. Review of Scientific Instruments 75, 2197–2210 (2004).

46. Jünger, F. et al. Measuring Local Viscosities near Plasma Membranes of Living Cells with Photonic Force Microscopy. Biophysical Journal 109, 869–882 (2015).

47. Gjorevski N. et al. Tissue geometry drives deterministic organoid patterning. Science 375, eaaw9021 (2022).

48. Schindelin, J. et al. Fiji: an open-source platform for biological-image analysis. Nature methods 9, 676–82 (2012).

49. Hörl, D. et al. BigStitcher: reconstructing high-resolution image datasets of cleared and expanded samples. Nat Methods 16, 870–874 (2019).

50. Wan, Y. et al. FluoRender: joint freehand segmentation and visualization for many-channel fluorescence data analysis. BMC Bioinformatics 18, 280 (2017).

50. Frisch, D. point2trimesh () - distance between point and triangulated surface. (https://www.mathworks.com/matlabcentral/fileexchange/52882-point2trimesh-distance-between-point-and-triangulated-surface), MATLAB Central File Exchange (2016). Retrieved December 1, 2021.

52. Dalgarno, P. A. et al. Multiplane imaging and three dimensional nanoscale particle tracking in biological microscopy. Optics Express 18, 877 (2010).

53. Edelstein, A., Amodaj, N., Hoover, K., Vale, R. & Stuurman, N. Computer Control of Microscopes Using μManager. Current Protocols in Molecular Biology 92, 14.20.1–14.20.17 (2010).

54. Tassieri, M., Evans, R. M. L., Warren, R. L., Bailey, N. J. & Cooper, J. M. Microrheology with optical tweezers: Data analysis. New Journal of Physics 14, 115032 (2012).

55. Smith, M. G., Gibson, G. M. & Tassieri, M. i-RheoFT: Fourier transforming sampled functions without artefacts. Sci Rep 11, 24047 (2021).

56. Matheson, A. B. et al. Microrheology With an Anisotropic Optical Trap. Front. Phys 9, 621512 (2021).

57. Ashworth, J. C. et al. Preparation of a User-Defined Peptide Gel for Controlled 3D Culture Models of Cancer and Disease. J Vis Exp e61710 (2020) doi:10.3791/61710.

